# Understanding genetic architecture overcomes tradeoffs between seed quality and insect resistance

**DOI:** 10.1101/2024.12.20.629859

**Authors:** Joseph R. White, James P. McNellie, Kyle G. Keepers, Brian C. Smart, Zoe M. Portlas, Zach E. Marcus, Nolan C. Kane, Jarrad R. Prasifka, Brent S. Hulke

## Abstract

The sunflower (*Helianthus annuus*) pericarp protects the seed within from both abiotic and biotic stresses. Achenes with stronger pericarps are less susceptible to damage from insect feeding. Complicating the genetic improvement of pericarp strength is the negative correlation between pericarp thickness (a component of strength) and oil content. As breeding efforts have increased oil content, there has been a concomitant decrease in pericarp thickness. A logical sunflower improvement goal is to improve oil content while preserving pericarp strength through genetic mechanisms independent of the tradeoffs with pericarp thickness. To determine the genetic basis of oil content, pericarp strength, and thickness, we identified QTL in two populations; the Sunflower Association Mapping panel (Mandel et al., 2011) and a recombinant inbred line (RIL) population derived from a thin pericarp oilseed inbred (HA 467) crossed to a thick pericarp open pollinated variety from Türkiye (PI 170415). A region on chromosome 15 was associated with neighboring QTL for banded moth resistance, oil content, and pericarp thickness, partially underlying the trade-offs among these traits. Additional QTL on chromosome 5 and 14 for pericarp strength provide fewer trade-offs with oil content. QTL for pericarp strength on chromosome 5 and pericarp thickness on chromosome 16 were associated with large structural variants, with candidate gene presence/absence variation between the haplotypes on chromosome 5. Understanding the origin and nature of phenotypic tradeoffs is beneficial to plant biologists and sunflower breeders as they seek to understand the origin and genetic architecture of adaptive and maladaptive traits.

## Introduction

Sunflower (*Helianthus annuus*) is grown for their achenes, a majority of which are processed to extract oil for human consumption. From 2000 to 2023, oilseed acreage in the United States averaged 1,512,525 acres (612,097 hectares) and confectionary sunflower comprised a further 282,323 acres (114,252 hectares; *NASS - Quick Stats*, 2023). Confectionery achenes are consumed directly by humans (i.e., not pressed for oil); the seeds are large and easy to dehull (i.e. to separate kernel from pericarp). Oil type sunflower achenes are small, black, difficult to dehull and contain over 40% oil (Petraru et al., 2021). Sunflower oil comprises 9% of the global vegetable oil market and is the 4^th^ most consumed seed oil behind rapeseed, palm, and soy (Economic Research Service, 2022). Selecting for high oil content has resulted in achenes with a thin pericarp and little to no unoccupied inner pericarp space. In other words, breeders have increased the kernel to pericarp ratio.

The pericarp is important during growth and development as it provides protection against insect pests. Several North American insect species have evolved as *Helianthus* specialists, namely the banded sunflower moth (*Cochylis hospes* Walsingham), sunflower head moth (*Homoeosoma electellum*), and red sunflower seed weevil (*Smicronyx fulvus*; Prasifka & Hulke, 2012). These insect pests penetrate the sunflower pericarp tissue when adults lay eggs (*S. fulvus*) or as larvae feed (*C. hospes* and *H. electellum*; Peng & Brewer, 1995), and severe infestations can result in substantial yield loss (Rogers, 1978). The geographic range and life cycle of these insects is affected by the environment, with milder winters associated with climate change often increasing their survival and distribution, thus expanding their potential economic damage (Barker & Enz, 1993; Debaeke et al., 2017; Pantzke et al., 2023; Prasifka, 2015; Royer & Knodel, 2019). Another obstacle for producers is reduction in labeled insecticides. Pesticides, such as chlorpyrifos, have high efficacy but deleterious effects on non-target organisms, including humans, and have been banned, reducing management options (Messina, 2021). A source of genetic resistance to *S. fulvus* has been identified but known resistance genes to *C. hospes* and *H. electellum* are lacking (DeGreef et al., 2020; Prasifka, 2020). Developing cultivars with improved insect resistance is a priority for sunflower researchers (Kleingartner, 2015). A better understanding of the genetic control of sunflower pericarp traits is part of a holistic strategy to enhance resistance against seed infesting insects.

The relationship between pericarp strength and *H. electellum* damage was previously discovered using bioassays that exposed developing achenes to larva in a laboratory setting. When given the choice, *H. electellum* larva preferentially consumed achenes with a weaker pericarp over achenes with a stronger pericarp (Prasifka et al., 2014; Sikora, 2017). Pericarp thickness has been hypothesized to contribute to pericarp strength; however, the role of pericarp thickness itself in resistance to *H. electellum* has not yet been established (Sikora, 2017). Identifying molecular markers associated with increased pericarp strength would facilitate breeding efforts towards cultivars with improved insect resistance. Mapping studies have been conducted for achene shape, but not for achene strength and thickness (Hasson et al., 2021; Reinert et al., 2020; Tang et al., 2006; Yue et al., 2009).

A challenge for sunflower breeders is to increase pericarp strength while maintaining a high oil concentration. An estimated 2/3 of the genetic gain in oil content can be attributed to decreased pericarp thickness and 1/3 to increased oil concentration in the kernel (Fick & Miller, 1997). Identifying loci that increase the strength of the pericarp while minimizing any decrease in oil content (i.e. increasing strength independent of thickness) would assist in creating cultivars that meet grower requirements.

The objective of this research is to gain a better understanding of the genetic control of pericarp properties and oil content to facilitate improvement of pericarp strength and insect resistance while minimizing reduction in oil content. To this end, we measured pericarp strength, pericarp thickness, oil content and damage from banded sunflower moth (*C. hospes*) in a biparental RIL population. Pericarp thickness and oil content were measured in the Sunflower Association Mapping (SAM) panel.

## Materials and Methods

### Population Development

#### Bi-parental RIL population

The two parents used to develop the RIL population, HA 467 (PI 670489) and PI 170415, possess different pericarp properties (Figure 1). HA 467 is an oilseed inbred line with a thin pericarp and high oleic acid released in 2006 by the USDA-ARS, Fargo, ND (Hulke et al., 2018). PI 170415 is an open pollinated variety with a thick pericarp obtained from Eceabat, Türkiye, in 1948, and deposited in the National Plant Germplasm System, USDA-ARS. HA 467 has a low mean pericarp strength of 2.62 N and PI 170415 a high mean pericarp strength of 4.49 N, measured on developing pericarps 14 days after anthesis. Furthermore, the cellular organization of the pericarp tissue differs between the two lines, with the higher pericarp strength associated with thicker pericarp and more organized cell walls (Figure 1). A single F1 progeny from the cross HA 467 × PI 170415 was self-pollinated and progeny were recurrently self-pollinated to the F7 generation to create 174 recombinant inbred lines.

**Figure 1:**
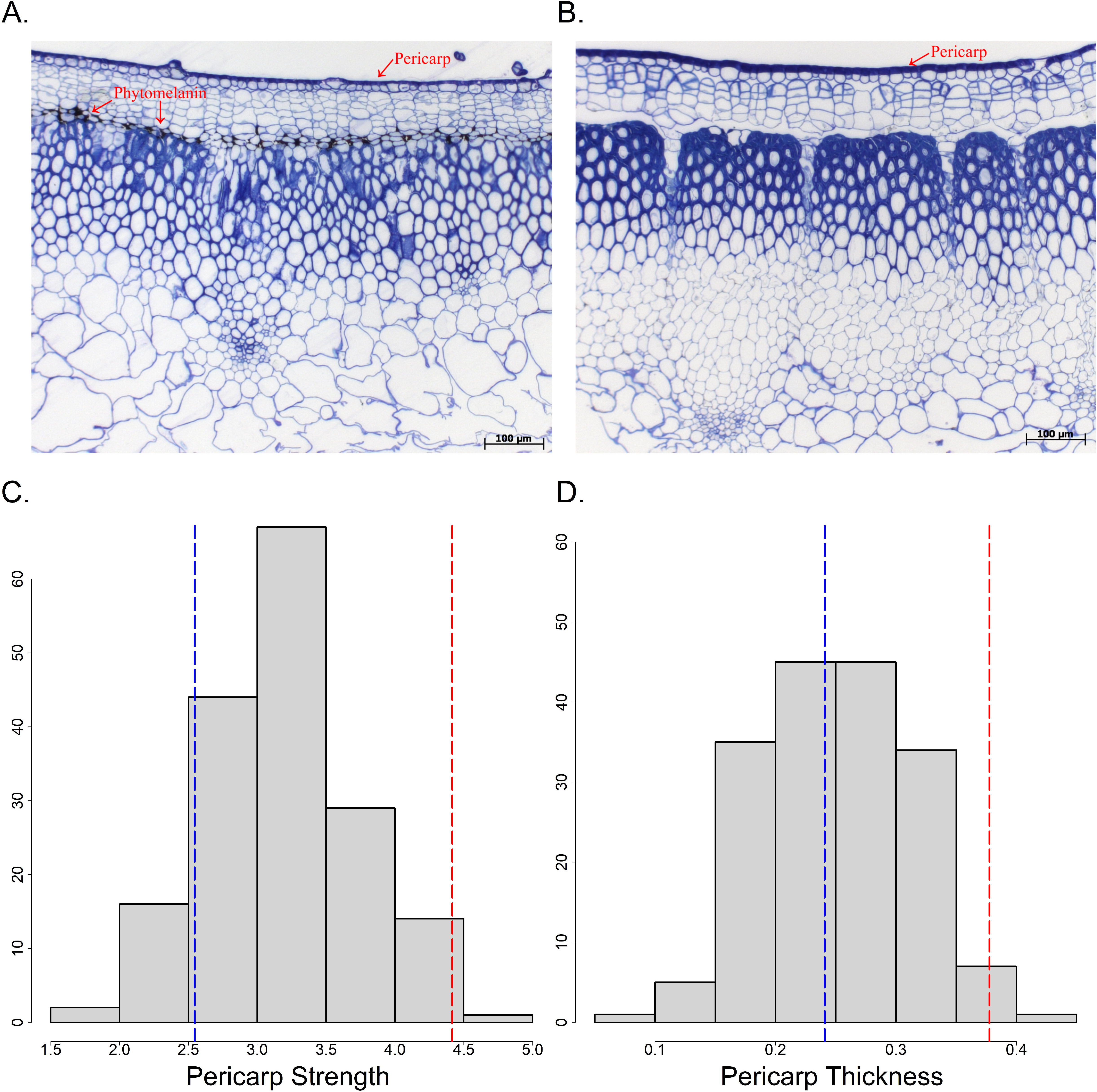
Light microscopy of mature achenes for HA 467 (A) and PI 170415 (B). Histogram of observed phenotypes for pericarp strength (C) and pericarp thickness (D) in the bi-parental recombinant inbred line population derived from HA 467 × PI 170415 grown in 2018 and 2022. Blue vertical lines denote observed values for HA 468 and red vertical lines observed values for PI 170415.

#### Sunflower Association Mapping Panel

The SAM panel is comprised of confectionery and oilseed inbred lines from the USDA and INRAE programs, improved accessions (i.e. open pollinated varieties), and landraces (Mandel et al., 2011). The SAM population has previously been used to map QTL for many phenotypes including carotenoids (Dowell & Mason, 2023), disease resistance (Yu et al., 2024), root morphology (Masalia et al., 2018), salinity tolerance (McNellie et al., 2024) and seed and floret size (Reinert et al., 2020).

### Phenotyping

#### Bi-parental RIL population

In 2018 and 2022, 174 F7 RILs were grown in Moorhead, MN, USA (46.87° N, 99.77° W). Fields were tilled and chemical weed control applied prior to emergence. Single row plots contained 25 plants, and were 6.10 m in length with 0.76 m between rows. Developing achenes were sampled at 14 days from the beginning of anthesis (R-5.1; Schneiter & Miller, 1981). This developmental stage was selected because it is the point where seed feeding larvae of both banded sunflower moth and sunflower head moth, transition from feeding on florets to developing achenes (Rogers, 1978). In other words, the sampling time is an approximation for when the pericarp is expected to be most important for protecting the seed from larval damage. Pericarp strength is the force, in Newtons (N), required to penetrate the hull of 14-day old achenes, measured using a Shimpo FGV-10XY Penetrometer with a No. 16 steel tapestry needle at the tip (Prasifka et al., 2014). For each genotype, penetrometer force was measured on five achenes for three heads and averaged.

Achenes from 3 to 5 plants per RIL were collected at maturity and used to measure pericarp thickness. Three achenes per plant were dissected and pericarp thickness measured using a digital caliper (Mitutoyo 547-500, Mitutoyo Corporation, Kawasaki, Japan). Because excised sections of pericarp (typically halves) were concave and the anvils of the calipers were flat, the tip of a sewing needle (American size 20) was affixed to one anvil to obtain accurate measurements. Data was collected from 159 RIL. An additional trait was derived by taking the residual of pericarp strength regressed on pericarp thickness, which we call strength independent of thickness (SIT). The residuals capture the proportion of pericarp strength not explained by pericarp thickness (Bian et al., 2014; Kelleher et al., 2018; Murray et al., 2008). The banded sunflower moth is a common insect pest in the field sites of this study, and frequently at economic levels of damage. Banded sunflower moth damage was calculated by taking X-ray images of 100 mature achenes. The resulting images were analyzed manually to determine the percentage of achenes with BSM damage. There were 125 genotypes with sufficient seed to measure oil content using an Oxford MQC NMR, fitted with a 51 mm probe. The instrument was calibrated using a collection of sunflower seed samples with varying oil content, quantified according to the Association of Official Agricultural Chemists (AOAC) method 2003.05 analysis which uses petroleum ether as the solvent (Minnesota Valley Testing Laboratories, New Ulm, MN, USA; J. Sieh, pers. comm.).

#### Sunflower Association Mapping Panel

Pericarp thickness and oil content was measured on 230 SAM accessions grown for 2 years (2015 and 2016) outside of Moorhead, MN. Single row plots contained 25 plants, and were 6.10 m in length with 0.76 m between rows. Flower heads were bagged and self-pollinated.

Pericarp thickness was measured on 2 to 3 achenes per head for 5 heads, as described for the bi-parental population. Oil content was measured as described for the bi-parental population.

### Genotyping

#### Bi-parental

Genomic DNA was extracted from leaf tissues using a Qiagen DNeasy 96 Plant Kit, quality validated using a Qubit DNA Concentration Assay, and resulting DNA stored at -20°C. DNA libraries were prepared using Twist 96-Plex Library Preparation Kits (Twist Bioscience, San Francisco, California, USA). Libraries were barcoded with i5 adapters and i7 index primers 11 and 12 from the kit, with a target insert size of 300 bp. Sequencing was conducted by Novogene (Sacramento, California, USA) and paired-end reads were generated with an average length of 151 bp. Genomic libraries were demultiplexed using the “*fgbio*” DemuxFastqs function with default parameters (Fennell & Homer, 2021). Quality control and processing of raw sequence data was conducted with FastQC and Fastp (Andrews, 2010; Chen, 2023; Chen et al., 2018). Trimmed reads were mapped to an indexed reference genome (HA 412 HOv.2; Miller et al., 2006) using BWA (Li, 2013) and further processed using samtools (McKenna et al., 2010). Variants were called against the reference genome on a per-sample basis using GATK HaplotypeCaller (Van der Auwera et al., 2013). Missing markers were imputed using Beagle v5.4 (Browning et al., 2018).

To explore the influence of a known inversion on the distal end of chromosome 5, the inverted region haplotype of each RIL had to be determined. This was accomplished in samtools by comparing the integral of read depths for regions inside and surrounding the inversion to two reference genomes (HA 412 HOv.2 and PSC8 [NCBI Accession GCA_026651735.1]), each possessing a different haplotype at the inversion site. The ratio of normalized read counts to the HA 412 HOv.2 genome within the inverted region was divided by normalized read counts in surrounding regions for each RIL. A depth ratio cutoff was chosen based on the difference between adjacent-ranked values, where a Mendelian segregation pattern would likely cause a change larger than expected from a ranked listing of stochastic depth integrals. Ratio values below 0.70 were considered to have the PSC8 inversion haplotype and values greater than or equal to 0.70 were considered to contain the HA 412 HO haplotype (Supplemental Figure 1). Read mapping was repeated with PSC8, with corroborating results. The ratio was then used to study the role of the inversion in pericarp strength via type III ANOVA and likelihood ratio test (Fox et al., 2012; R Team, 2023).

#### Sunflower Association Mapping Panel

The SAM panel was previously re-sequenced at a 10x average depth and aligned to the HA 412 HOv.2 reference genome (Hübner et al., 2019; Todesco et al., 2020). Single nucleotide polymorphisms (SNP) were filtered to remove variants below a minor allele frequency of 0.05 and/or greater than 40 heterozygous calls. Linkage disequilibrium pruning was performed using PLINK (*--indep-pairwise 50 5 0.95*) to remove redundant markers (Purcell et al., 2007). There were 223,907 SNP remaining for use in subsequent QTL mapping.

### Data analysis and QTL mapping

#### Bi-parental

Least squared genotype means for pericarp strength, thickness, SIT, and oil content were calculated in ASReml-R (Butler et al., 2017) using the following model:

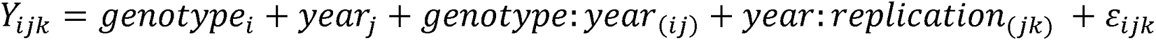

Where *Y_ijk_* is the phenotype of an observational unit, *genotype_i_*, the effect of the *i*th genotype; *year_j_*, the effect of the *j*th year; genotype: *year*_(*ij*)_, the interaction of the *i*th genotype with the *j*th year; year: replication_(*jk*)_, the effect of the *k*th replication in the *j*th year and *ε_ijk_* is the residual error. Genotype and year were considered fixed effects and the other effects were random. Variance components were estimated using the same model for the purpose of calculating heritability, but with genotype as a random effect. The coefficient of variation (CV, the square root of the error variance divided by the mean multiplied by 100), and variance components were calculated in R using ASreml-R. The equation for broad sense heritability on an entry-mean basis (*h*^2^) is:

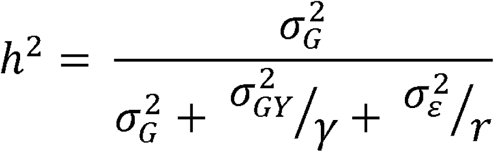

where genetic variance is denoted as 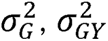 is the genotype × year interaction variance, 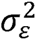 is the residual variance, *γ* the harmonic mean of number of years per genotype, and *r* is the harmonic mean of number of plots per genotype (Holland et al., 2003). Pearson correlation coefficients were calculated in R (R Team, 2023).

SNP were recoded so that QTL effect can be attributed to a parental allele; AA for HA 467 and BB for PI 170415. The genetic map was constructed in R/qtl and contained 2,397 markers, spanning 3,567 cM (Broman et al., 2003). QTL were discovered using inclusive composite interval mapping in IciMapping (Li et al., 2007). Significance thresholds were calculated via a permutation test with 1000 iterations. The functional identity of putative genes within 2 LOD confidence intervals were inferred using the Eugene v.1.1 annotation of the HA 412 HOv.2 Sunflower Genome Brower (sunflowergenome.org) JBrowse2 (Diesh et al., 2023) and National Center for Biotechnology Information - National Library of Medicine (NCBI-NLM) Basic Local Alignment Search Tool (BLAST; Altschul et al., 1990). Genes within the region of low homology in the inversion on chromosome 5 were identified using reciprocal protein BLAST searches (blastp) between the candidate genes to the *Arabidopsis thaliana* annotated genome (NCBI accession <u>GCA_000001735.1</u>). The sunflower protein sequence used was obtained from https://sunflowergenome.org/annotations-data/.

#### Sunflower Association Mapping Panel

Least squared means for oil content and pericarp thickness were calculated using the MIXED procedure in SAS (SAS Institute, 2018) using the same model as in the bi-parental population. Coefficient of variation, correlation, variance analysis and heritability were calculated as described in the bi-parental population. Association between phenotype and SNP was conducted in GEMMA using a Wald significance test (Zhou & Stephens, 2012, 2014). Principal components were used to control for population structure as fixed effect and genomic relationship matrix to control for relatedness as a random effect. The principal component analysis and a genomic relationship matrix were calculated using PLINK (Purcell et al., 2007). Models were fit using 0 to 8 principal components; after post-hoc inspection of QQ and Manhattan plots, models with 3 principal components were selected. The logarithm of the odds (LOD) threshold for QTL significance was 6.65, determined using a Bonferroni correction of 0.05 divided by the number of SNP (223,907). That threshold was relaxed to 6.60 (equivalent to a significance threshold of p=0.056 before Bonferroni correction) to include an additional QTL for pericarp thickness on chromosome 12. Putative genes were identified following the same procedure as with the bi-parental population but within 100,000 bp upstream and downstream of significant SNP.

## Results

### Bi-parental

Pericarp strength and thickness were measured on 159 RIL derived from HA 467, a maintainer line with thin pericarp and a high oil content, crossed with PI 170415, a thick pericarp landrace from Türkiye. Pericarp strength had a mean of 3.15 N, ranging from 1.89 to 4.54 N, and a coefficient of variation of 14.62 (Figure 1, Table 1). Pericarp thickness had a mean of 0.24 mm, ranging from 0.10 to 0.40 mm, and a CV of 17.82. Oil content had a mean of 237 g oil per kg seed mass, ranging from 106 to 439, and a CV of 9.88. Percent banded sunflower moth (BSM) damage to achenes had a mean of 20.6 %, ranging from 4.8 % to 54.0 % and a CV of 53.39. Entry mean basis, broad sense heritability estimates ranged from 0.53 for banded sunflower moth damage to 0.88 for oil content. Pericarp strength and thickness had a Pearson correlation coefficient of 0.50, suggesting that pericarp thickness only explains 25 % of the phenotypic variation in pericarp strength. The correlation between oil content and SIT (-0.21, p < 0.05) was lower compared to pericarp strength (-0.54, p < 0.0001) and pericarp thickness (-0.67, p < 0.0001; Table 2).

**Table 1:** Mean, variance components and heritability for 5 traits measured in 159 recombinant inbred lines derived from HA 467 × PI 170415 and two traits measured in 230 accessions of the sunflower association mapping population.

| Recombinant Inbred Lines |  |  |  |  |  |  |
| --- | --- | --- | --- | --- | --- | --- |
| Trait | Mean | CV<sup>a</sup> | $\sigma^2_g$ | $\sigma^2_{gy}$ | $\sigma^2_\varepsilon$ | $h^2$ |
| Pericarp Strength | 3.15 N | 14.62 | 0.25 (46.14%) | 0.07 (13.69%) | 0.21 (39.35%) | 0.76 |
| Pericarp Thickness | 0.24mm | 17.82 | 0.0032 (55.71%) | 0.0007 (11.37%) | 0.0019 (32.92%) | 0.84 |
| SIT<sup>b</sup> | 0.00 | 0.00 | 0.18 (52.54%) | 0.02 (6.82%) | 0.14 (40.06%) | 0.84 |
| Oil Content | 236.53 g seed kg<sup>-1</sup> seed mass | 9.88 | 2273.41 (80.62%) | -<sup>e</sup> | 546.37 (19.38%) | 0.88 |
| BSM<sup>c</sup> Damage | 20.60% | 53.39 | 92.08 (30.17%) | 92.11 (30.18%) | 120.99 (39.65%) | 0.53 |
| Sunflower Association Mapping |  |  |  |  |  |  |
| Trait | Mean | CV<sup>a</sup> | $\sigma^2_g$ | $\sigma^2_{gy}$ | $\sigma^2_\varepsilon$ | $h^2$ |
| Oil Content | 335.76 | 11.58 | 5491.44 (71%) | 821.84 (11%) | 1381.63 (18%) | 0.87 |
| Pericarp Thickness | 0.23 | 16.91 | 0.0112 (83%) | 0.0008 (6%) | 0.0015 (11%) | 0.93 |

<sup>a</sup> CV, coefficient of variation
<sup>b</sup> SIT, strength independent of thickness
<sup>c</sup> BSM, Banded sunflower moth
<sup>e</sup> Oil content was evaluated in a single year

**Table 2:**
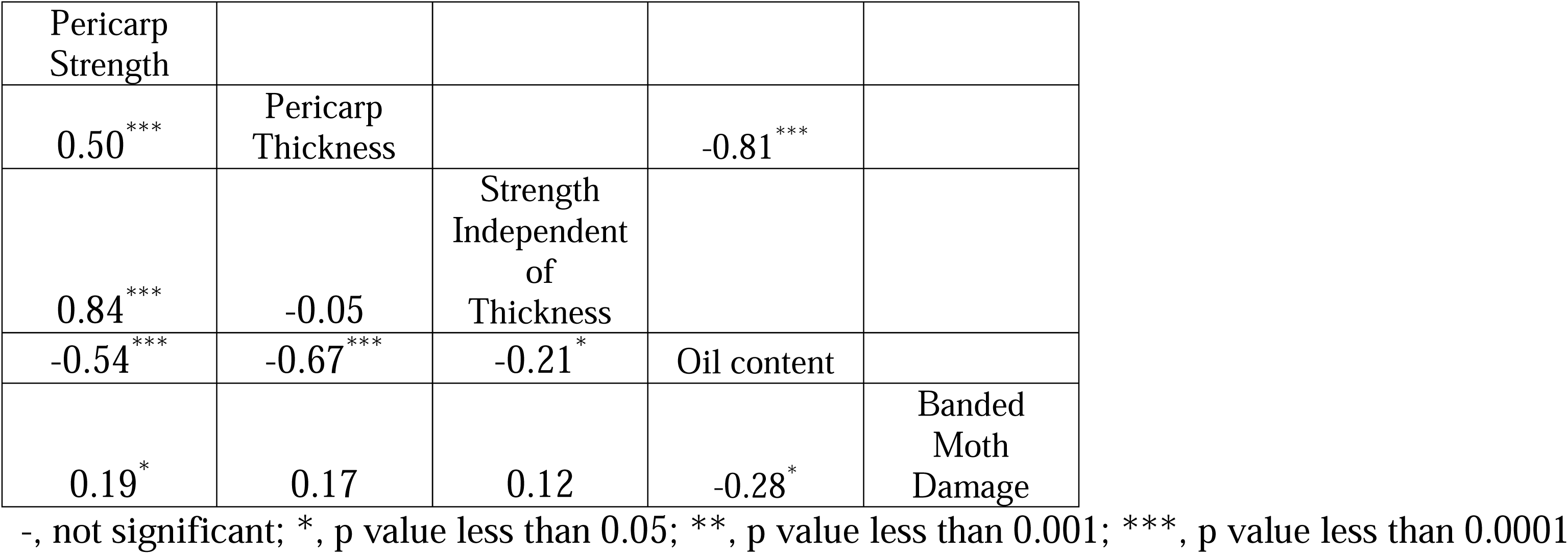
Pearson correlation coefficient for 5 traits measured in the recombinant inbred lines derived from HA 467 × PI 170415 (lower triangle) and for two traits measured in the sunflower association mapping population (upper triangle).

#### QTL Mapping

Twelve significant QTL were mapped in the bi-parental population, two each for pericarp strength, pericarp strength independent of thickness, and BSM damage; five for pericarp thickness; and one for oil content (Figure 2; Table 3). For pericarp strength, QTL were located on chromosomes 5 (between 62,522,539 and 62,626,373 bp), and 14 (between 21,728,213 and 21,772,687 bp) explaining 12.68% and 9.19% of variation respectively. For pericarp thickness, QTL were mapped to chromosomes 2 (between 176,970,777 and 177,057,599 bp), 4 (between 207,204,709 and 207,328,329 bp), 15 (between 61,508,948 and 61,985,314 bp) and 16 (between 193,951,480 and 196,960,718 bp for the first QTL, and for the second QTL, between 216,467,659 and 216,525,958 bp). Phenotypic variation explained was 7.15%, 8.06%, and 11.50% for QTL on chromosomes 2, 4, and 15, while the two QTL on chromosome 16 explained 7.21% and 8.44% of phenotypic variation. Of the alleles inherited from HA 467, QTL on chromosomes 4, 15, and 16 decreased pericarp thickness, as expected, while the QTL on chromosome 2 increased pericarp thickness. SIT resulted in two QTL, the first on chromosome 5 (17.34% variation explained) at the same position as the QTL for pericarp strength. The second QTL for SIT mapped to chromosome 14 between 79,423,855 and 79,937,827bp, 58 Mbp from the QTL on chromosome 14 for pericarp strength. There is evidence that the QTL on chromosome 14 are genetically linked. The linkage map shows limited recombination between QTL (Supplemental Figure 2) and QTL effects are similar (Table 4). The single QTL for oil content mapped to chromosome 15 between 58,617,255 and 58,864,941 bp explaining 11.57% of variation, with the allele from HA 467 conferring an increase in oil. BSM damage had two QTL, the first on chromosome 13 (between 24,188,305 and 25,346,013 bp) explaining 12.01% of variation and another near the pericarp thickness and oil content QTL on chromosome 15 (between 68,101,718 and 70,008,248 bp) explaining 8.92% of variation. The allele from HA 467 increased damage (i.e. decreased resistance) on chromosome 13 and decreased damage (improved resistance) on chromosome 15. There were 169 predicted genes within the 2 LOD drop QTL interval, 125 of which had predicted gene products (Supplemental Table 1).

**Figure 2:**
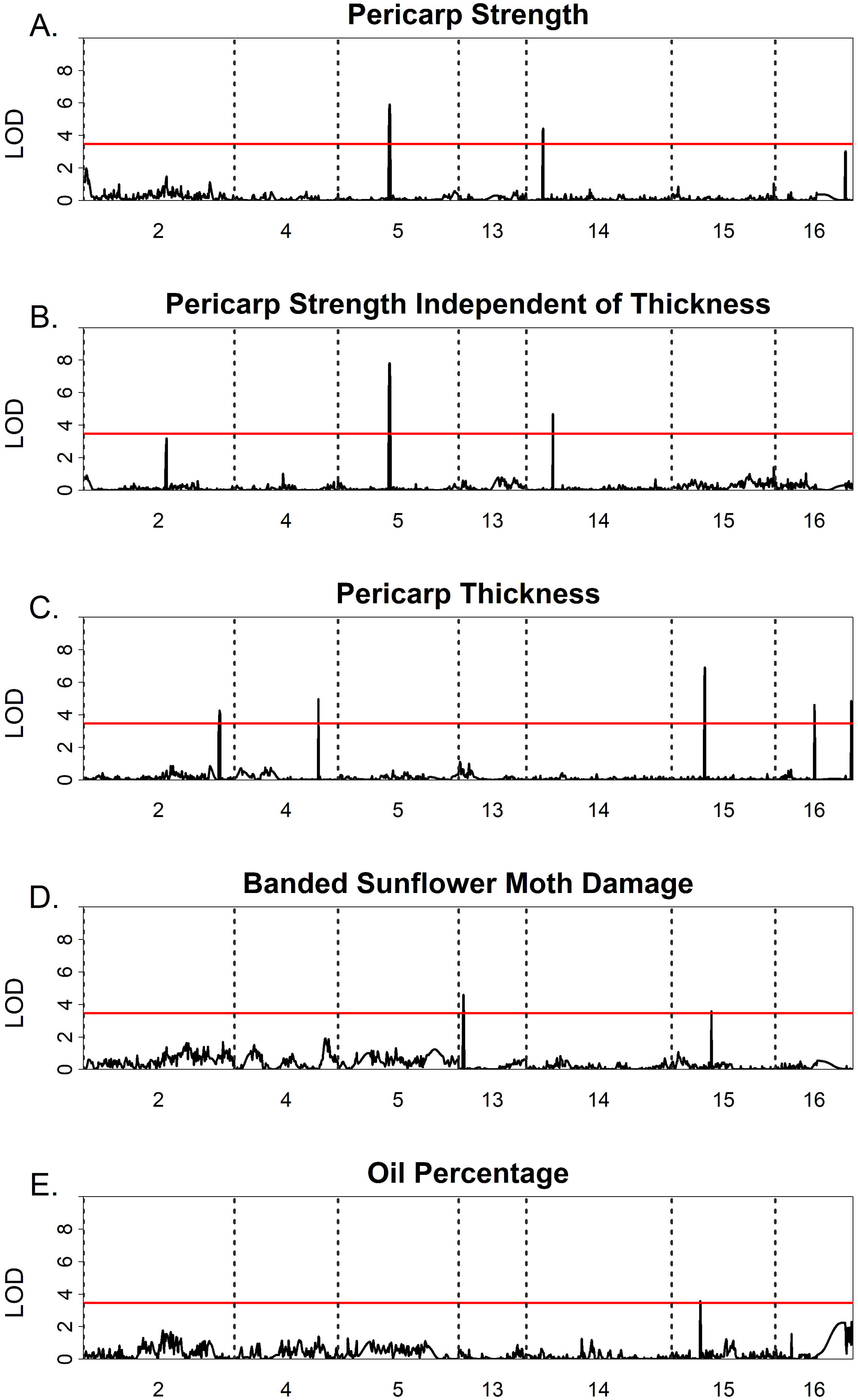
Genetic mapping results using 159 recombinant inbred lines derived from HA 467 × PI 170415 for pericarp strength (A), pericarp strength independent of thickness (B), and pericarp thickness (C). Horizontal red lines denote significance threshold calculated using 1000 permutations.

**Table 3.** QTL mapping results in bi-parental recombinant inbred lines (top) and in the sunflower association mapping panel (bottom). QTL for pericarp thickness were found in both populations, while traits with QTL unique to the bi-parental population are pericarp strength, strength independent of thickness (SIT) and banded sunflower moth (BSM) damage. Oil content QTL reported in the sunflower association mapping population.

| Trait | Chromosome | Left Marker | Right Marker | PVE<sup>a</sup> | LOD | Effect<sup>c</sup> |
| --- | --- | --- | --- | --- | --- | --- |
| Pericarp Strength | 5 | 62,522,539 | 62,626,373 | 12.68% | 5.91 | 0.19 |
|  | 14 | 21,728,213 | 21,772,687 | 9.19% | 4.43 | -0.16 |
| Pericarp Thickness | 2 | 176,970,777 | 177,057,599 | 7.15% | 4.27 | 0.02 |
|  | 4 | 207,204,709 | 207,328,329 | 8.06% | 4.97 | -0.02 |
|  | 15 | 61,508,948 | 61,985,314 | 11.50% | 6.92 | -0.02 |
|  | 16 | 193,951,480 | 196,960,718 | 7.21% | 4.61 | -0.02 |
|  | 16 | 216,467,659 | 216,525,958 | 8.44% | 4.85 | -0.02 |
| SIT<sup>b</sup> | 5 | 62,522,539 | 62,626,373 | 17.34% | 7.82 | 0.19 |
|  | 14 | 79,423,855 | 79,937,827 | 9.41% | 4.68 | -0.14 |
| Oil content | 15 | 58,617,255 | 58,864,941 | 11.57% | 3.59 | 1.77 |
| BSM Percent Damage | 13 | 24,188,305 | 25,346,013 | 12.01% | 4.60 | 2.05 |
|  | 15 | 68,101,718 | 70,008,248 | 8.92% | 3.59 | -1.77 |

Sunflower Association Mapping Panel
| Trait | Chromosome | Position<sup>d</sup> | Reference Allele | Alternative Allele | LOD | Effect<sup>c</sup> |
| --- | --- | --- | --- | --- | --- | --- |
| Oil content | 15 | 144,019,238 | C | T | 7.08 | -1.82 |
| Pericarp Thickness | 7 | 153,182,010 | C | T | 7.22 | 0.06 |
|  | 10 | 111,456,652 | T | C | 6.97 | 0.07 |
|  | 10 | 122,754,494 | T | C | 7.16 | 0.06 |
|  | 12 | 9,810,492 | A | T | 6.64 | 0.05 |
|  | 16 | 198,639,052 | G | A | 7.16 | 0.04 |

<sup>a</sup> PVE, Percent variation explained
<sup>b</sup> SIT, strength independent of thickness
<sup>c</sup> The effect of the allele donated from HA 467 in the bi-parental RIL population. In the sunflower association mapping panel, the QTL effect is the effect of substituting the reference genome allele for the alternative allele.
<sup>d</sup> QTL position based upon reference genome HA 412 HO

**Table 4:** Additive QTL effect for all traits measured in **a** recombinant inbred lines derived from HA 467 × PI 170415. Each row corresponds to a QTL with the underlined value denoting which trait the QTL was significant.

| Chromosome | Position <sup>a</sup> | Pericarp strength | Pericarp thickness | SIT<sup>b</sup> | Banded sunflower moth damage | Oil content |
| --- | --- | --- | --- | --- | --- | --- |
| 2 | 176,970,777 | 0.041<sup>c</sup> | <u>0.016</u> | 0.007 | -0.792 | -0.312 |
| 4 | 207,204,709 | -0.040 | <u>-0.017</u> | 0.002 | -0.056 | 1.041 |
| 5 | 62,522,539 | <u>0.189</u> | -0.001 | <u>0.189</u> | 0.741 | -0.863 |
| 13 | 24,188,305 | 0.006 | -0.006 | 0.042 | <u>2.054</u> | 0.511 |
| 14 | 21,728,213 | <u>-0.160</u> | -0.000 | -0.023 | 0.214 | 0.558 |
| 14 | 79,423,855 | -0.002 | -0.001 | <u>-0.138</u> | 0.056 | 0.191 |
| 15 | 58,617,255 | -0.022 | -0.002 | 0.030 | <u>-1.768</u> | -0.192 |
| 15 | 61,508,948 | -0.030 | <u>-0.020</u> | 0.031 | 0.218 | -0.101 |
| 15 | 68,101,718 | -0.022 | -0.003 | 0.032 | 0.038 | <u>1.767</u> |
| 16 | 193,951,480 | -0.030 | <u>-0.016</u> | 0.018 | -0.474 | 0.269 |
| 16 | 216,467,659 | -0.002 | <u>-0.017</u> | -0.040 | -0.029 | 0.852 |

Sunflower Association Mapping Panel
| Chromosome | Position | Pericarp strength | Pericarp thickness | SIT | Banded sunflower moth damage | Oil content |
| --- | --- | --- | --- | --- | --- | --- |
| 7 | 144,019,238 | - | <u>0.06</u> | - | - | 0.07 |
| 10 | 111,456,652 | - | <u>0.06</u> | - | - | -2.00 |
| 10 | 153,182,010 | - | <u>0.07</u> | - | - | -0.86 |
| 12 | 122,754,494 | - | <u>0.05</u> | - | - | -0.62 |
| 15 | 198,639,052 | - | 0.03 | - | - | <u>-1.82</u> |
| 16 | 9,810,492 | - | <u>0.04</u> | - | - | -1.22 |

<sup>a</sup> QTL position based upon reference genome HA 412 HO. For the bi-parental population QTL position is the physical position of the left closest marker.
<sup>b</sup> SIT, strength independent of thickness
<sup>c</sup> The effect of the allele donated from HA 467 in the bi-parental RIL population. In the sunflower association mapping panel, the QTL effect is the effect of substituting the reference genome allele for the alternative allele.

The three QTL on chromosome 15 are genetically linked based on the genetic map and few observed recombination events (Supplemental Figure 3); however, QTL effect estimates at the three loci across traits suggest independent genetic control, sometimes with opposing effects (Table 4). The HA 467 (i.e. thin pericarp) allele of the thickness QTL also increases BSM damage, as expected, but is in phase with an allele at the nearby BSM damage QTL that decreases BSM damage. Interestingly, a high oil content allele at the oil QTL is in phase with thin pericarp, low oil allele at the thickness QTL, suggesting that the tradeoff between thickness and oil content is driven by linkage, at least on chromosome 15 (Table 4).

There was an inversion detected near QTL on chromosome 5 (pericarp strength and SIT; Supplemental Figure 4) and a structural variant between the QTL for pericarp thickness on chromosome 16 (Supplemental Figure 5). The prior work of Todesco et al. (2020) informed us that the chromosome 5 haplotypes are both common in cultivated germplasm. An analysis of read depth indicates that HA 467 contains a haplotype very similar to the HA 412 HO haplotype and PI 170415 contains one similar to the PSC8 haplotype. The haplotype from HA 467 has a negative correlation with pericarp strength (r = -0.36) and 112 RIL contained this haplotype versus 47 that did not (Supplemental Figure 1). The apparent segregation distortion at this site could contribute to this inverted region not being identified in linkage mapping analyses. To test if the inversion explained pericarp strength better than the detected QTL on chromosome 5, a linear model was fit containing pericarp strength as the response variable and the QTL on chromosome 5 and the read depth ratio as independent variables. Type III ANOVA determined that the inversion is better at explaining pericarp variation than the QTL on chromosome 5. This was confirmed by a likelihood ratio test that found a significant improvement to model fit when the inversion is added to the model (Supplemental Table 2). Genes within each inversion haplotype were annotated and their physical position compared (Supplemental Figure 4). Three distinct regions emerged: two inversions with relatively high sequence homology between reference genomes separated by a 5.8Mb region in the PSC8 genome that is absent in the HA 412 HO genome. Within that 5.8Mb region in PSC8, there are 17 copies of *Hydroxycinnamoyl-CoA shikimate transferase* (*HCT*). *HCT* is present only once in the larger of the two inversions in the HA 412 HO (HA 467) haplotype (Supplemental Table 3). Two of the 17 copies are over 90% identical to the HA 412 HO *HCT* homolog, with the other 15 copies 54 to 70% identical. Also present within the region are 3 full copies of the transcription factor *ULTRAPETALA* and 4 copies of *ABC transporter G family member 9.* Both genes are present only once in the HA 412 HO haplotype. Long read de novo assembly data for the parental haplotypes is not available for the structural variation on chromosome 16; however, similar presence/absence variation could be present there.

#### Sunflower Association Mapping Population

The SAM population is comprised of two market classes (oilseed and confectionery) and two heterotic groups. The two heterotic groups are denoted as maintainers (B) and restorers (R), referring to the cytoplasmic male sterility system. For both market classes, the R heterotic group had a lower mean for pericarp thickness and a higher mean for oil content (Supplemental Table 4). When all market classes and heterotic groups were analyzed together, the CV for oil content was 11.58 and 16.91 for pericarp thickness. Mean oil content was 336 g oil kg^-1^ seed mass, ranging from 65 to 512. Mean pericarp thickness was 0.23 mm, ranging from 0.04 to 0.74. Heritability estimates were high, 0.87 for oil content, and 0.93 for pericarp thickness (Table 1). As in the bi-parental population, oil content was negatively correlated with pericarp thickness (r = -0.81; Table 2).

#### Genome-wide Association

Oil content mapped as a single QTL on chromosome 15 at 144,019,238 bp, with the HA 412 HO reference allele conferring a increase of 1.82% (Table 4). There were five significant QTL for pericarp thickness, located on chromosome 7 at 153,182,010 bp, chromosome 10 at 111,456,652 bp and 122,754,494 bp, chromosome 12 at 9,810,492 bp, and chromosome 16 at 198,639,052 bp. The QTL for pericarp thickness on chromosome 16 is between the two QTL detected in the RIL population, suggesting that the QTL from the SAM population is located within the inversion detected in the bi-parental population. The HA 412 HO reference allele conferred an decrease in pericarp thickness for all significant SNP, ranging from 0.04 mm on chromosome 16 to 0.07 mm on chromosome 10 at 111,456,652 bp. There are 39 predicted genes within 100,000 bp of QTL peaks, 23 of which had predicted gene products (Supplemental Table 1).

## Discussion

Our research identified the genetic basis of pericarp and seed properties, including traits with significant, unwanted correlations resulting in challenging trade-offs for plant breeders. Specifically, we examined the genetic control of pericarp strength, thickness, pericarp strength independent of thickness, oil content, and banded sunflower moth damage/resistance with the objective of identifying alleles that can improve pericarp strength and oil content while avoiding their negative correlation. We found genetic loci underlying the trade-offs between pericarp strength and oil content, as well as loci that affect variation in individual traits with little or no contribution to trade-offs. Previous research has shown that sunflower head moth larvae avoided achenes with a strong pericarp, suggesting that improved resistance of immature achenes to larvae predation could be obtained by selecting for a strong pericarp (Prasifka et al., 2014). In the field environments of this study, we did not see sunflower head moth in large enough numbers to provide a useful phenotype. Instead, we studied damage due to a similar lepidopteran pest, the banded sunflower moth, which occurs in our region. Our work provided the first report of QTL associated with BSM damage and these loci deserve further study in additional populations. Within the two BSM QTL, 66 putative genes with predicted protein products were found. On chromosome 13, Ha412HOChr13g0587101 (Ha412HOChr12:25066852..25071678) encodes a thioredoxin domain containing protein, which is part of the reactive oxygen species response to insect damage in *Arabidopsis thaliana* and *Brachypodium distachyon* (Collins et al., 2010; Subramanyam et al., 2019). It is promising that genes with putative insect response functions in other plant species are within QTL intervals for BSM damage.

A clear genetic signal indicating that banded sunflower moth damage is reduced by increased pericarp strength at 14 d post-anthesis is not apparent at first glance. The HA 467 allele at the QTL for BSM damage on chromosome 15 confers a decrease in damage but is linked to a QTL for decreased pericarp thickness (Supplementary Figure 3, Table 4). This contributes to the unexpected positive correlation between BSM damage and pericarp thickness (Table 2). The HA 467 allele of another QTL for pericarp thickness (chromosome 2) confers a significant increase in pericarp thickness and a sizeable reduction in BSM damage.

The relationship between oil content and pericarp thickness in the SAM panel roughly follows the pattern reported in 1947; that for a 1% reduction in the pericarp, oil increases by 0.47-0.75% (Radanović et al., 2018). In the SAM panel, the reference allele for the QTL for pericarp thickness on chromosome 15 at 144,019,238 bp confers an estimated 12.60% decrease in pericarp thickness and an increase of 1.81% in oil content (Table 4). This QTL was previously identified in the SAM population with a different phenotypic data set at 145,002,138 bp (McNellie et al., 2024). While thickness does enhance strength, 75% of the phenotypic variance in strength is independent of thickness, suggesting that high oil and high strength pericarp can occur together. The negative correlation between oil content and SIT was less than the correlation between oil content and pericarp strength and thickness. This makes increasing SIT a good breeding goal for multi-trait selection strategies involving oil content because SIT limits negative trade-offs. Even so, we established that the one significant QTL for oil content in the bi-parental population was only negatively associated with pericarp thickness by linkage. This oil QTL is also associated with a candidate gene that is not expected to have an effect on pericarp traits: a putative 3-ketoacyl-CoA synthase gene, required for elongation of fatty acids, about 160 kbp away (Blacklock & Jaworski, 2006; James Jr et al., 1995). Two other copies of *H. annuus* 3-ketoacyl-CoA synthase were previously cloned and characterized but are located on chromosomes 13 and 16 (González-Mellado et al., 2019).

There was one instance of QTL co-localizing between populations, for pericarp thickness on chromosome 16. The two QTL on chromosome 16 for pericarp thickness in the RIL population flank a structural variant, possibly a translocation (Supplemental Figure 5). The QTL for pericarp thickness in the SAM population mapped between the physical position of the 2 QTL in the RIL population. No QTL for pericarp properties in the current study co-localized with previously reported QTL for other achene traits. There are 38 predicted genes within QTL intervals for pericarp strength and 27 within the QTL intervals for pericarp thickness.

The location of QTL for pericarp strength and SIT on chromosome 5 is near a known inversion (ann05.01; Todesco et al., 2020). This inversion was recently found to have a role in oleic acid stability in inbred lines (Markus Ingold, pers. Comm., 2023). The alleles of each parental haplotype were found to be common in the public germplasm and important for other traits, as well (Todesco et al., 2020; Huang et al., 2023). Our report of a novel region containing multiple copies of *HCT*, *ULTRAPETALA*, and an ABC transporter within the PSC8 haplotype of this region may explain the correlation of this inversion with pericarp strength. *HCT* has an established role in lignin production (Bonawitz & Chapple, 2010; Hoffmann et al., 2004; Shu et al., 2021). Lignin is present in sunflower achenes, especially in the sclerenchyma, and contributes to achene strength (Hernández & Bellés, 2007). We hypothesize that the extra copies of *HCT* in the PSC8 inversion result in an increase in lignin deposition in the pericarp. *ULTRAPETALA* has a known role in floral meristem processes and epigenetic complexes affecting chromatin remodeling and histone modification (Moreau et al., 2016; Ornelas-Ayala et al., 2021), and may be involved with pericarp development, as the pericarp is a maternally inherited floral tissue. ABC transporters are involved in a plethora of genetic processes, including lignin transportation and seed size (Alejandro et al., 2012; Fabre et al., 2016; Theodoulou & Kerr, 2015). While their involvement with pericarp traits is hypothetical, the presence of multiple copies in this unique region within the chromosome 5 inversion raises the question of their role in pericarp strength. Functional genomics experiments with these genes with respect to lignin deposition are needed to confirm their roles in pericarp structure.

Our discoveries suggest (1) that most of the variability in pericarp strength is independent of thickness, (2) that some of the negative correlation of thickness with oil content is the result of genetic linkage, and (3) additional variation in seed infesting insect resistance is available, independent of pericarp strength. Tradeoffs that are due to closely linked loci will likely persist in breeding populations unless marker-assisted tools derived from these findings are used to develop novel haplotypes, particularly on chromosome 15. While some variation for these traits is due to large structural variants, these tend to contribute very little to tradeoffs. This suggests that the goal of developing cultivars with a strong pericarp and high oil content is possible. More broadly, implication of structural variants in pericarp traits suggest interesting biology in the evolution of the pericarp. Our findings add to the growing body of literature that important genetic variation may be concentrated in large inversions, potentially derived from introgression across species boundaries. Obtaining a better understanding of where structural variations reside within the *Helianthus annuus* genome and the effect of structural variation on traits of interest is needed.

## Abbreviations

BSM: banded sunflower moth
CV: coefficient of variation
HCT: Hydroxycinnamoyl-CoA shikimate transferase
QTL: quantitative trait loci
RIL: recombinant inbred lines
SAM: sunflower association mapping
SIT: strength independent of thickness

## Statements and Declarations

## Funding

The authors wish to acknowledge funding from U.S. Department of Agriculture – Agricultural Research Service CRIS project 3060-21000-043-00D and 3060-21000-047-00D. McNellie is supported by a U.S. Department of Agriculture – Agricultural Research Service Postdoctoral Fellowship. This work used resources of the Center for Computationally Assisted Science and Technology (CCAST) at North Dakota State University, which were made possible in part by NSF MRI Award No. 2019077.

### Acknowledgements

The authors gratefully acknowledge the technical efforts of Brady Koehler and the many interns of the USDA-ARS Hulke Lab.

## Data Availability

Data will be made available upon acceptance in the National Agricultural Library – Ag Data Commons repository. Code is available from GitHub at https://github.com/BrianSmart/

## Author Contributions

JRP and BSH conceived of the research. JRP, BSH, and NCK provided resources and supervision. JRW and JRP collected field data. JRW, JPM, ZMP, and ZEM collected laboratory data. JPM, JRW, BCS and KGK performed statistical analysis. JPM and JRW wrote the manuscript with input from other authors. All authors have read and approved the final manuscript.

## References

Alejandro, S., Lee, Y., Tohge, T., Sudre, D., Osorio, S., Park, J.,…Fernie, A. R. (2012). AtABCG29 is a monolignol transporter involved in lignin biosynthesis. Current Biology, 22(13), 1207–1212.

Altschul, S. F., Gish, W., Miller, W., Myers, E. W., & Lipman, D. J. (1990). Basic local alignment search tool. Journal of molecular biology, 215(3), 403–410.

Andrews, S. (2010). FastQC: a quality control tool for high throughput sequence data. In: Cambridge, United Kingdom.

Barker, J., & Enz, J. (1993). Development of laboratory reared banded sunflower moth, Cochylis hospes Walsingham (Lepidoptera: Cochylidae), in relation to temperature.

Bian, Y., Yang, Q., Balint-Kurti, P. J., Wisser, R. J., & Holland, J. B. (2014). Limits on the reproducibility of marker associations with southern leaf blight resistance in the maize nested association mapping population. BMC genomics, 15, 1–15.

Blacklock, B. J., & Jaworski, J. G. (2006). Substrate specificity of Arabidopsis 3-ketoacyl-CoA synthases. Biochemical and biophysical research communications, 346(2), 583–590.

Bonawitz, N. D., & Chapple, C. (2010). The genetics of lignin biosynthesis: connecting genotype to phenotype. Annual review of genetics, 44(1), 337–363.

Broman, K. W., Wu, H., Sen, Ś., & Churchill, G. A. (2003). R/qtl: QTL mapping in experimental crosses. bioinformatics, 19(7), 889–890.

Browning, B. L., Zhou, Y., & Browning, S. R. (2018). A one-penny imputed genome from next-generation reference panels. The American Journal of Human Genetics, 103(3), 338–348.

Butler, D., Cullis, B., Gilmour, A., Gogel, B., & Thompson, R. (2017). ASReml-R reference manual version 4. VSN International Ltd, Hemel Hempstead, HP1 1ES, UK.

Chen, S. (2023). Ultrafast one-pass FASTQ data preprocessing, quality control, and deduplication using fastp. Imeta, 2(2), e107.

Chen, S., Zhou, Y., Chen, Y., & Gu, J. (2018). fastp: an ultra-fast all-in-one FASTQ preprocessor. Bioinformatics, 34(17), i884–i890.

Collins, R. M., Afzal, M., Ward, D. A., Prescott, M. C., Sait, S. M., Rees, H. H., & Tomsett, A. B. (2010). Differential proteomic analysis of Arabidopsis thaliana genotypes exhibiting resistance or susceptibility to the insect herbivore, Plutella xylostella. PLoS One, 5(4), e10103.

Debaeke, P., Casadebaig, P., Flenet, F., & Langlade, N. (2017). Sunflower crop and climate change: vulnerability, adaptation, and mitigation potential from case-studies in Europe. OCL Oilseeds and fats crops and lipids, 24(1), 15 p.

DeGreef, M. G., Prasifka, J. R., Koehler, B. D., & Hulke, B. S. (2020). Registration of oilseed sunflower maintainer germplasm HA 488, with resistance to the red sunflower seed weevil. Journal of Plant Registrations, 14(2), 203–205.

Diesh, C., Stevens, G. J., Xie, P., De Jesus Martinez, T., Hershberg, E. A., Leung, A.,…Bridge, C. (2023). JBrowse 2: a modular genome browser with views of synteny and structural variation. Genome biology, 24(1), 1–21.

Dowell, J. A., & Mason, C. (2023). Candidate pathway association and genome-wide association approaches reveal alternative genetic architectures of carotenoid content in cultivated sunflower (Helianthus annuus). Applications in Plant Sciences, 11(6), e11558.

Economic Research Service, U. S. D. o. A. (2022). Oil Crop Outlook, April 2022. https://www.ers.usda.gov/data-products/chart-gallery/gallery/chart-detail/?chartId=104023

Fabre, G., Garroum, I., Mazurek, S., Daraspe, J., Mucciolo, A., Sankar, M.,…Nawrath, C. (2016). The ABCG transporter PEC1/ABCG32 is required for the formation of the developing leaf cuticle in Arabidopsis. New Phytologist, 209(1), 192–201.

Fennell, T., & Homer, N. (2021). fgbio: A set of tools to analyze genomic data with a focus on Next Generation Sequencing. In https://github.com/fulcrumgenomics/fgbio?tab=readmeov-file

Fick, G. N., & Miller, J. F. (1997). Sunflower breeding. Sunflower technology and production, 35, 395–439.

Fox, J., Weisberg, S., Adler, D., Bates, D., Baud-Bovy, G., Ellison, S.,…Graves, S. (2012). Package ‘car’. Vienna: R Foundation for Statistical Computing, 16(332), 333.

González-Mellado, D., Salas, J. J., Venegas-Calerón, M., Moreno-Pérez, A. J., Garcés, R., & Martínez-Force, E. (2019). Functional characterization and structural modelling of Helianthus annuus (sunflower) ketoacyl-CoA synthases and their role in seed oil composition. Planta, 249, 1823–1836.

Hasson, H., Dudhe, M., Mandel, T., Warschefsky, E., Rieseberg, L., & Hübner, S. (2021). Image processing and genome-wide association studies in sunflower identify loci associated with seed-coat characteristics. bioRxiv, 04.

Hernández, L., & Bellés, P. (2007). A 3-D finite element analysis of the sunflower (Helianthus annuus L.) fruit. Biomechanical approach for the improvement of its hullability. Journal of Food Engineering, 78(3), 861–869.

Hoffmann, L., Besseau, S., Geoffroy, P., Ritzenthaler, C., Meyer, D., Lapierre, C.,…Legrand, M. (2004). Silencing of hydroxycinnamoyl-coenzyme A shikimate/quinate hydroxycinnamoyltransferase affects phenylpropanoid biosynthesis. The Plant Cell, 16(6), 1446–1465.

Holland, J. B., Nyquist, W. E., Cervantes-Martínez, C. T., & Janick, J. (2003). Estimating and interpreting heritability for plant breeding: an update. Plant breeding reviews, 22.

Huang, K., Jahani, M. Gouzy, J., and Rieseberg, L.H. 2023. The genomics of linkage drag in inbred lines of sunflower. PNAS 120:e2205783119.

Hulke, B., Ma, G., Qi, L., & Gulya, T. (2018). Registration of oilseed sunflower germplasms RHA 461, RHA 462, RHA 463, HA 465, HA 466, HA 467, and RHA 468 with diversity in Sclerotinia resistance, yield, and other traits. Journal of Plant Registrations, 12(1), 142–147.

Institute, S. (2018). SAS 9.4 language reference: Concepts. SAS Institute.

James Jr, D. W., Lim, E., Keller, J., Plooy, I., Ralston, E., & Dooner, H. K. (1995). Directed tagging of the Arabidopsis FATTY ACID ELONGATION1 (FAE1) gene with the maize transposon activator. The Plant Cell, 7(3), 309–319.

Kelleher, E. S., Jaweria, J., Akoma, U., Ortega, L., & Tang, W. (2018). QTL mapping of natural variation reveals that the developmental regulator bruno reduces tolerance to P-element transposition in the Drosophila female germline. PLoS biology, 16(10), e2006040.

Kleingartner, L. (2015). US and Canada perspectives on sunflower production and processing. In Sunflower (pp. 491-516). Elsevier.

Li, H. (2013). Aligning sequence reads, clone sequences and assembly contigs with BWA-MEM. arXiv preprint arXiv:1303.3997.

Li, H., Ye, G., & Wang, J. (2007). A modified algorithm for the improvement of composite interval mapping. Genetics, 175(1), 361–374.

Mandel, J., Dechaine, J. M., Marek, L., & Burke, J. (2011). Genetic diversity and population structure in cultivated sunflower and a comparison to its wild progenitor, Helianthus annuus L. Theoretical and Applied Genetics, 123, 693–704.

Masalia, R. R., Temme, A. A., Torralba, N. d. l., & Burke, J. M. (2018). Multiple genomic regions influence root morphology and seedling growth in cultivated sunflower (Helianthus annuus L.) under well-watered and water-limited conditions. PLoS One, 13(9), e0204279.

McKenna, A., Hanna, M., Banks, E., Sivachenko, A., Cibulskis, K., Kernytsky, A.,…Daly, M. (2010). The Genome Analysis Toolkit: a MapReduce framework for analyzing next-generation DNA sequencing data. Genome research, 20(9), 1297–1303.

McNellie, J. P., May, W. E., Rieseberg, L. H., & Hulke, B. S. (2024). Association studies of salinity tolerance in sunflower provide robust breeding and selection strategies under climate change. Theoretical and Applied Genetics, 137(8), 184.

Messina, E. (2021). Chlorpyrifos Tolerance Revocations. Environmental Protection Agency.

Miller, J., Gulya, T., & Vick, B. (2006). Registration of three maintainer (HA 456, HA 457, and HA 412 HO) high-oleic oilseed sunflower germplasms. Crop science, 46(6), 2728.

Moreau, F., Thévenon, E., Blanvillain, R., Lopez-Vidriero, I., Franco-Zorrilla, J. M., Dumas, R.,…Carles, C. C. (2016). The Myb-domain protein ULTRAPETALA1 INTERACTING FACTOR 1 controls floral meristem activities in Arabidopsis. Development, 143(7), 1108–1119.

Murray, S. C., Sharma, A., Rooney, W. L., Klein, P. E., Mullet, J. E., Mitchell, S. E., & Kresovich, S. (2008). Genetic improvement of sorghum as a biofuel feedstock: I. QTL for stem sugar and grain nonstructural carbohydrates. Crop Science, 48(6), 2165–2179.

NASS - Quick Stats ((2023). USDA National Agricultural Statistics Service.

Ornelas-Ayala, D., Garay-Arroyo, A., García-Ponce, B., R. Álvarez-Buylla, E., & Sanchez, M. d. l. P. (2021). The epigenetic faces of ULTRAPETALA1. Frontiers in Plant Science, 12, 637244.

Pantzke, S., Ferguson, B., Rajamohan, A., Rinehart, J. P., Prischmann-Voldseth, D., & Prasifka, J. R. (2023). Thermal biology and overwintering behavior of the red sunflower seed weevil (Coleoptera: Curculionidae). Environmental Entomology, 52(4), 632–638.

Peng, C., & Brewer, G. J. (1995). Description of achene damage by the red sunflower seed weevil, the banded sunflower moth, and the sunflower moth. Journal of the Kansas Entomological Society, 68(3), 263–267.

Petraru, A., Ursachi, F., & Amariei, S. (2021). Nutritional characteristics assessment of sunflower seeds, oil and cake. Perspective of using sunflower oilcakes as a functional ingredient. Plants, 10(11), 2487.

Prasifka, J. R. (2015). Sunflower Insect Pests. Sunflower: Chemistry, Production, Processing, and Utilization, 157.

Prasifka, J. R. (2020). Susceptibility of sunflower inbred lines and putative resistance sources to Smicronyx fulvus LeConte (Coleoptera: Curculionidae). Journal of Applied Entomology, 144(7), 632–636.

Prasifka, J. R., & Hulke, B. S. (2012). Current status and future perspectives on sunflower insect pests. 18th International Sunflower Conference Program and Abstracts, Mar del Plata and Balcarce, Argentina,

Prasifka, J. R., Hulke, B. S., & Seiler, G. J. (2014). Pericarp strength of sunflower and its value for plant defense against the sunflower moth, Homoeosoma electellum. Arthropod-Plant Interactions, 8, 101–107.

Purcell, S., Neale, B., Todd-Brown, K., Thomas, L., Ferreira, M. A., Bender, D.,…Daly, M. J. (2007). PLINK: a tool set for whole-genome association and population-based linkage analyses. The American journal of human genetics, 81(3), 559–575.

Radanović, A., Miladinović, D., Cvejić, S., Jocković, M., & Jocić, S. (2018). Sunflower genetics from ancestors to modern hybrids—A review. Genes, 9(11), 528.

Reinert, S., Gao, Q., Ferguson, B., Portlas, Z. M., Prasifka, J. R., & Hulke, B. S. (2020). Seed and floret size parameters of sunflower are determined by partially overlapping sets of quantitative trait loci with epistatic interactions. Molecular Genetics and Genomics, 295, 143–154.

Rogers, C. E. (1978). Sunflower moth: feeding behavior of the larva. Environmental Entomology, 7(5), 763–765.

Royer, T. A., & Knodel, J. J. (2019). Sunflower Moth (Lepidoptera: Pyralidae) Biology, Ecology, and Management. Journal of Integrated Pest Management, 10(1), 1w–1w.

Schneiter, A., & Miller, J. (1981). Description of sunflower growth stages 1. Crop science, 21(6), 901–903.

Shu, F., Jiang, B., Yuan, Y., Li, M., Wu, W., Jin, Y., & Xiao, H. (2021). Biological activities and emerging roles of lignin and lignin-based products1 A review. Biomacromolecules, 22(12), 4905–4918.

Sikora, D. M. (2017). Evaluation of host plant resistance against sunflower moth, Homoeosoma electellum (Hulst), in cultivated sunflower in western Nebraska.

Subramanyam, S., Nemacheck, J. A., Hargarten, A. M., Sardesai, N., Schemerhorn, B. J., & Williams, C. E. (2019). Multiple molecular defense strategies in Brachypodium distachyon surmount Hessian fly (Mayetiola destructor) larvae-induced susceptibility for plant survival. Scientific Reports, 9(1), 2596.

Tang, S., Leon, A., Bridges, W. C., & Knapp, S. J. (2006). Quantitative trait loci for genetically correlated seed traits are tightly linked to branching and pericarp pigment loci in sunflower. Crop Science, 46(2), 721–734.

Team, R. C. (2023). R: A Language and Environment for Statistical Computing. In R Foundation for Statistical Computing. https://www.R-project.org/

Theodoulou, F. L., & Kerr, I. D. (2015). ABC transporter research: going strong 40 years on. Biochemical Society Transactions, 43(5), 1033–1040.

Todesco, M., Owens, G. L., Bercovich, N., Légaré, J.-S., Soudi, S., Burge, D. O.,…Imerovski, I. (2020). Massive haplotypes underlie ecotypic differentiation in sunflowers. Nature, 584(7822), 602–607.

Van der Auwera, G. A., Carneiro, M. O., Hartl, C., Poplin, R., Del Angel, G., Levy-Moonshine, A.,…Thibault, J. (2013). From FastQ data to high-confidence variant calls: the genome analysis toolkit best practices pipeline. Current protocols in bioinformatics, 43(1), 11.10. 11–11.10. 33.

Yu, Y., Yang, J., Zhang, J., Rieseberg, L. H., & Zhao, J. (2024). Genomic Insights into Disease Resistance in Sunflower (Helianthus annuus): Identifying Key Regions and Candidate Genes for Verticillium dahliae Resistance. Plants, 13(18), 2582.

Yue, B., Cai, X., Yuan, W., Vick, B., & Hu, J. (2009). Mapping quantitative trait loci (QTL) controlling seed morphology and disk diameter in sunflower (Helianthus annuus L.). Helia, 32(50), 17–35.

Zhou, X., & Stephens, M. (2012). Genome-wide efficient mixed-model analysis for association studies. Nature genetics, 44(7), 821–824.

Zhou, X., & Stephens, M. (2014). Efficient multivariate linear mixed model algorithms for genome-wide association studies. Nature methods, 11(4), 407–409.

