## Supplemental Figure for "Understanding genetic architecture overcomes tradeoffs between seed quality and insect resistance"

### Slide 1
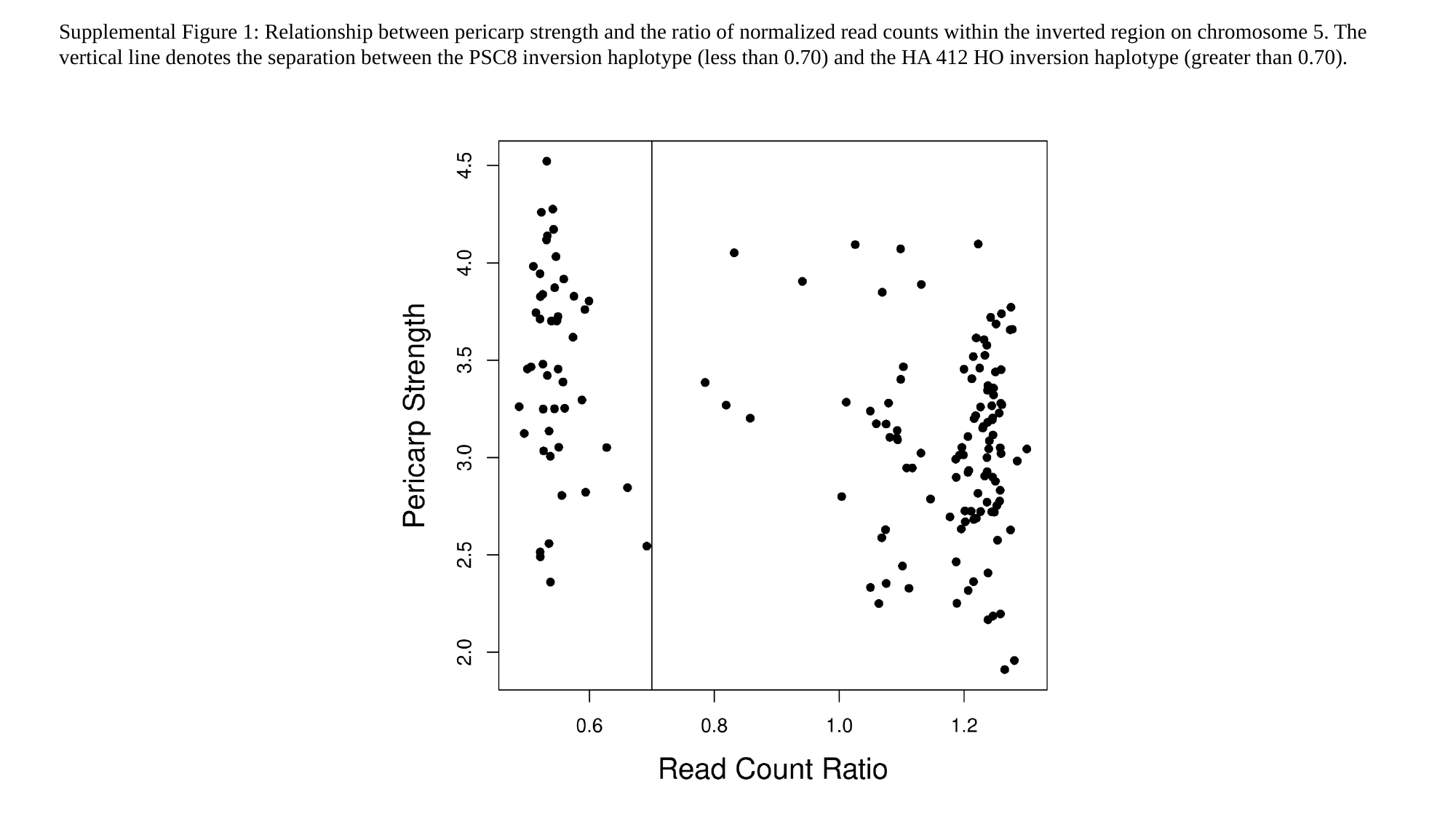

Supplemental Figure 1: Relationship between pericarp strength and the ratio of normalized read counts within the inverted region on chromosome 5. The vertical line denotes the separation between the PSC8 inversion haplotype (less than 0.70) and the HA 412 HO inversion haplotype (greater than 0.70).

### Slide 2
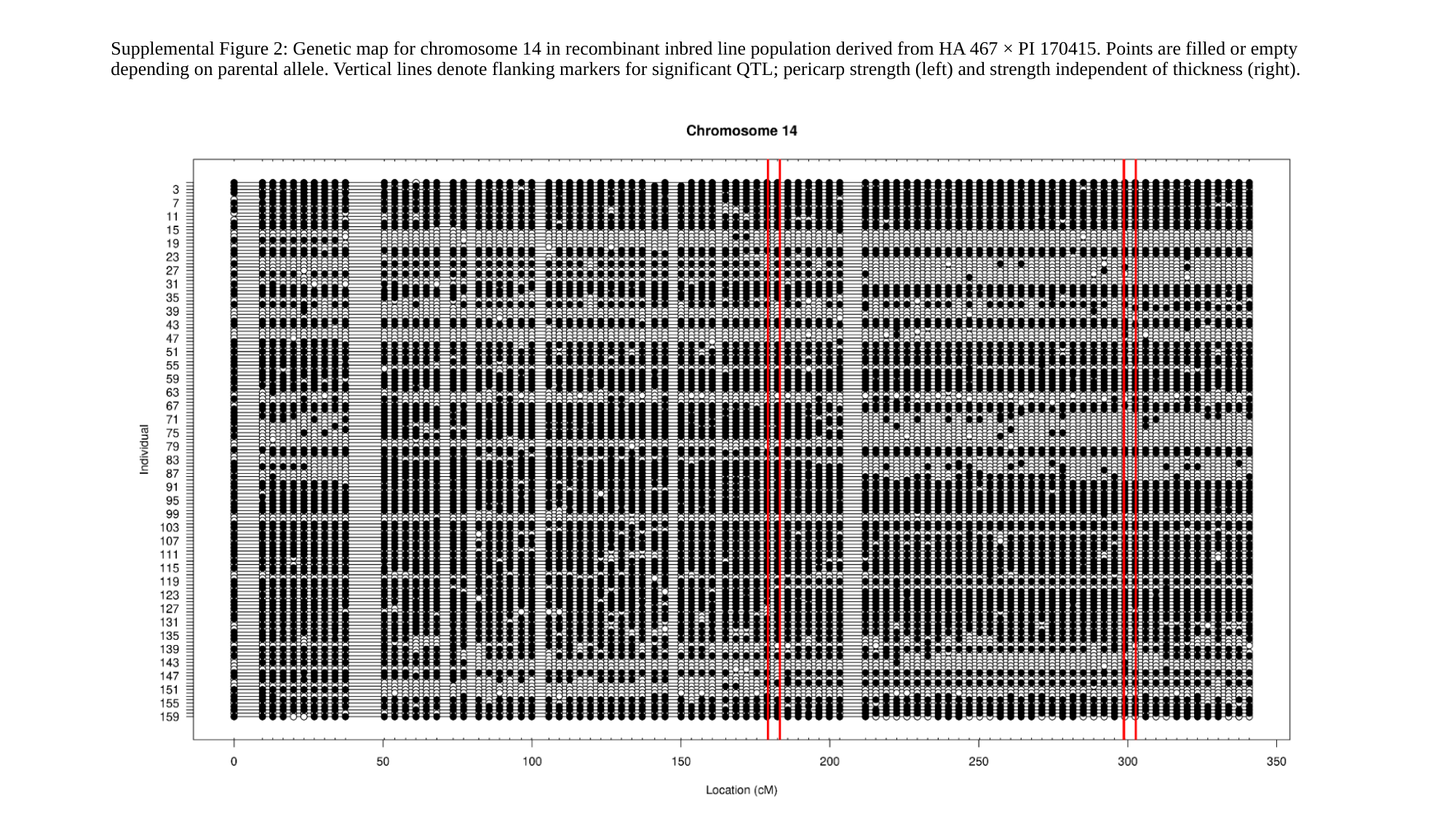

#### Supplemental Figure 2: Genetic map for chromosome 14 in recombinant inbred line population derived from HA 467 × PI 170415. Points are filled or empty depending on parental allele. Vertical lines denote flanking markers for significant QTL; pericarp strength (left) and strength independent of thickness (right).

### Slide 3
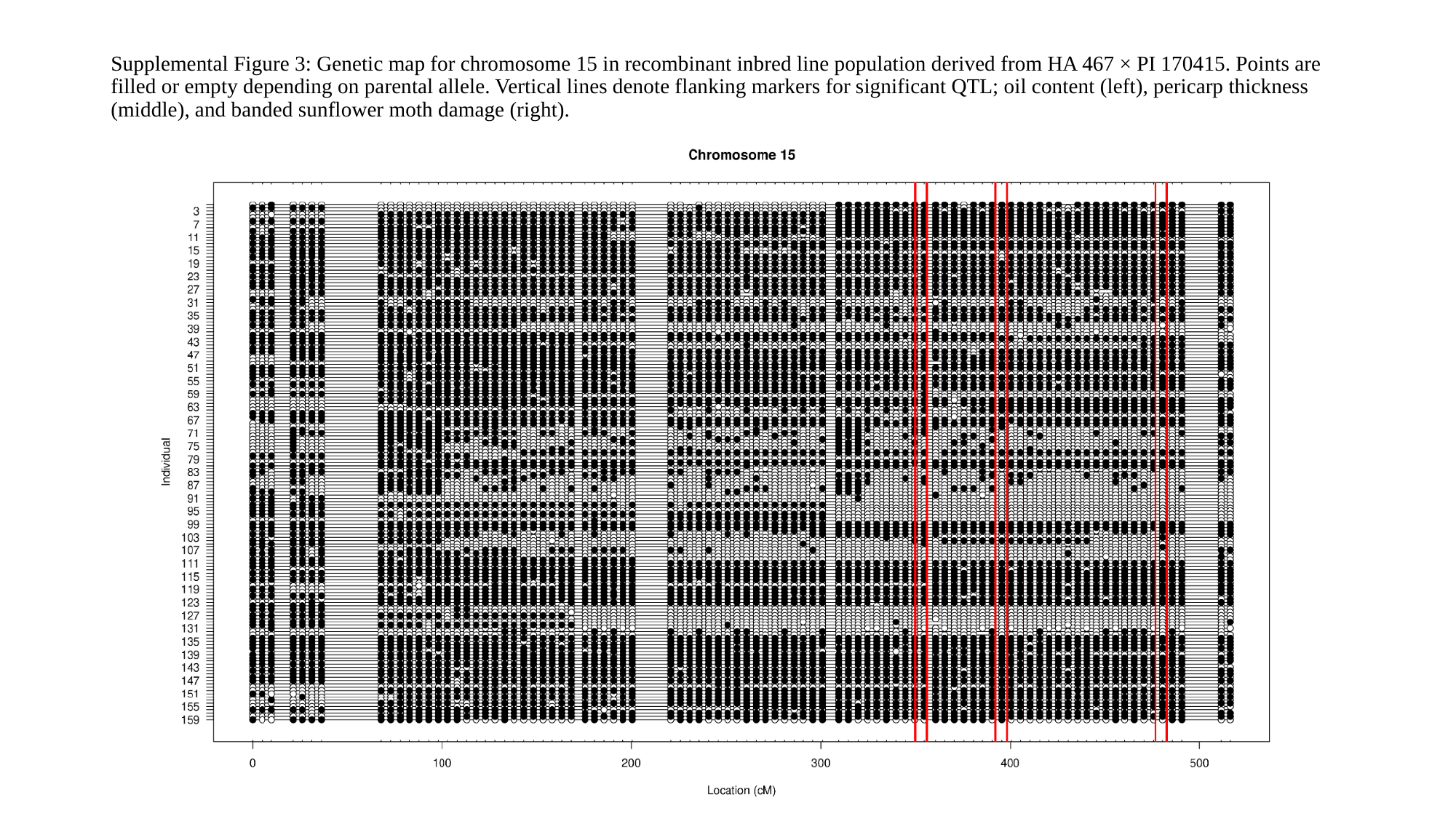

#### Supplemental Figure 3: Genetic map for chromosome 15 in recombinant inbred line population derived from HA 467 × PI 170415. Points are filled or empty depending on parental allele. Vertical lines denote flanking markers for significant QTL; oil content (left), pericarp thickness (middle), and banded sunflower moth damage (right).

### Slide 4
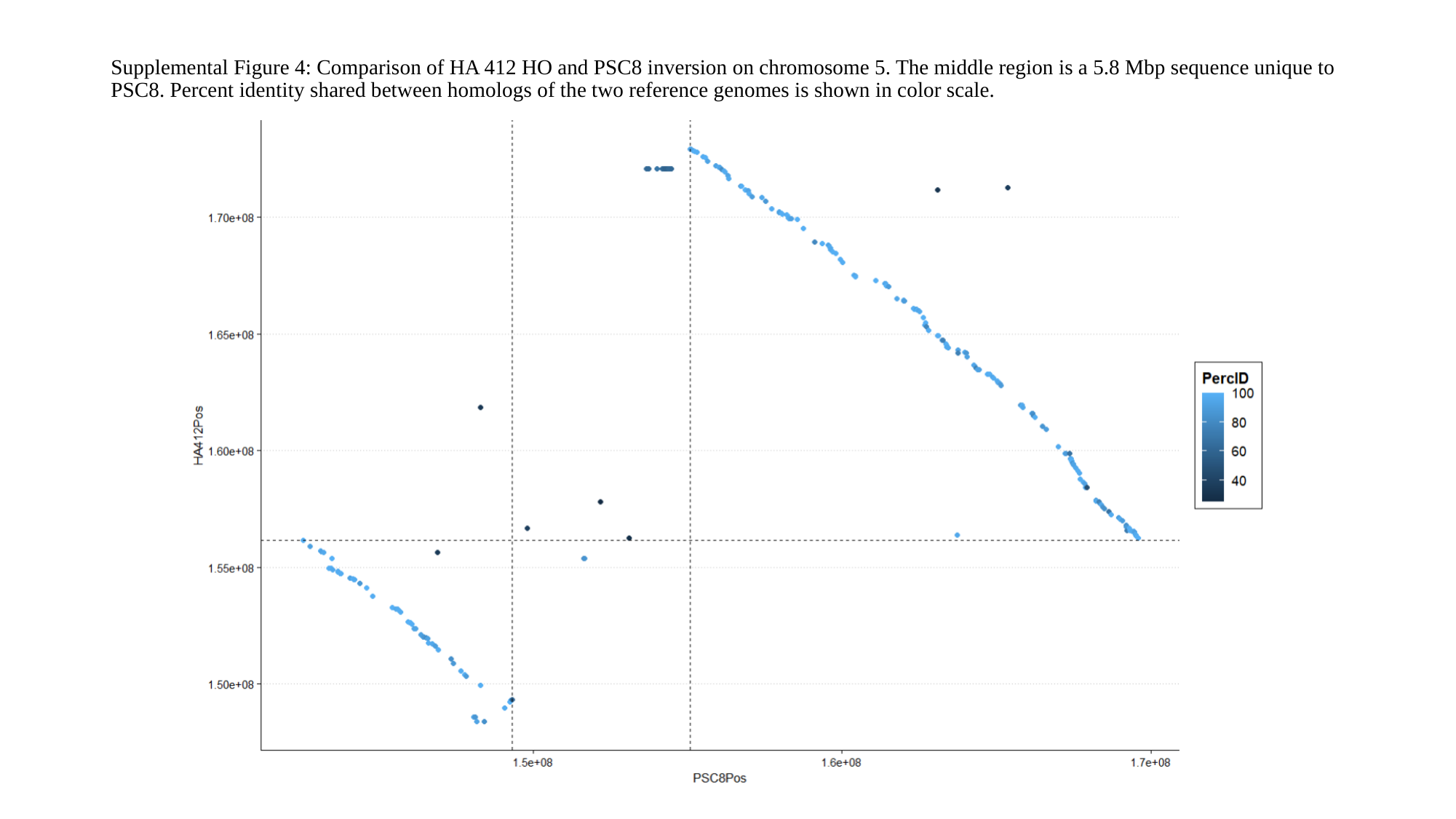

#### Supplemental Figure 4: Comparison of HA 412 HO and PSC8 inversion on chromosome 5. The middle region is a 5.8 Mbp sequence unique to PSC8. Percent identity shared between homologs of the two reference genomes is shown in color scale.

### Slide 5
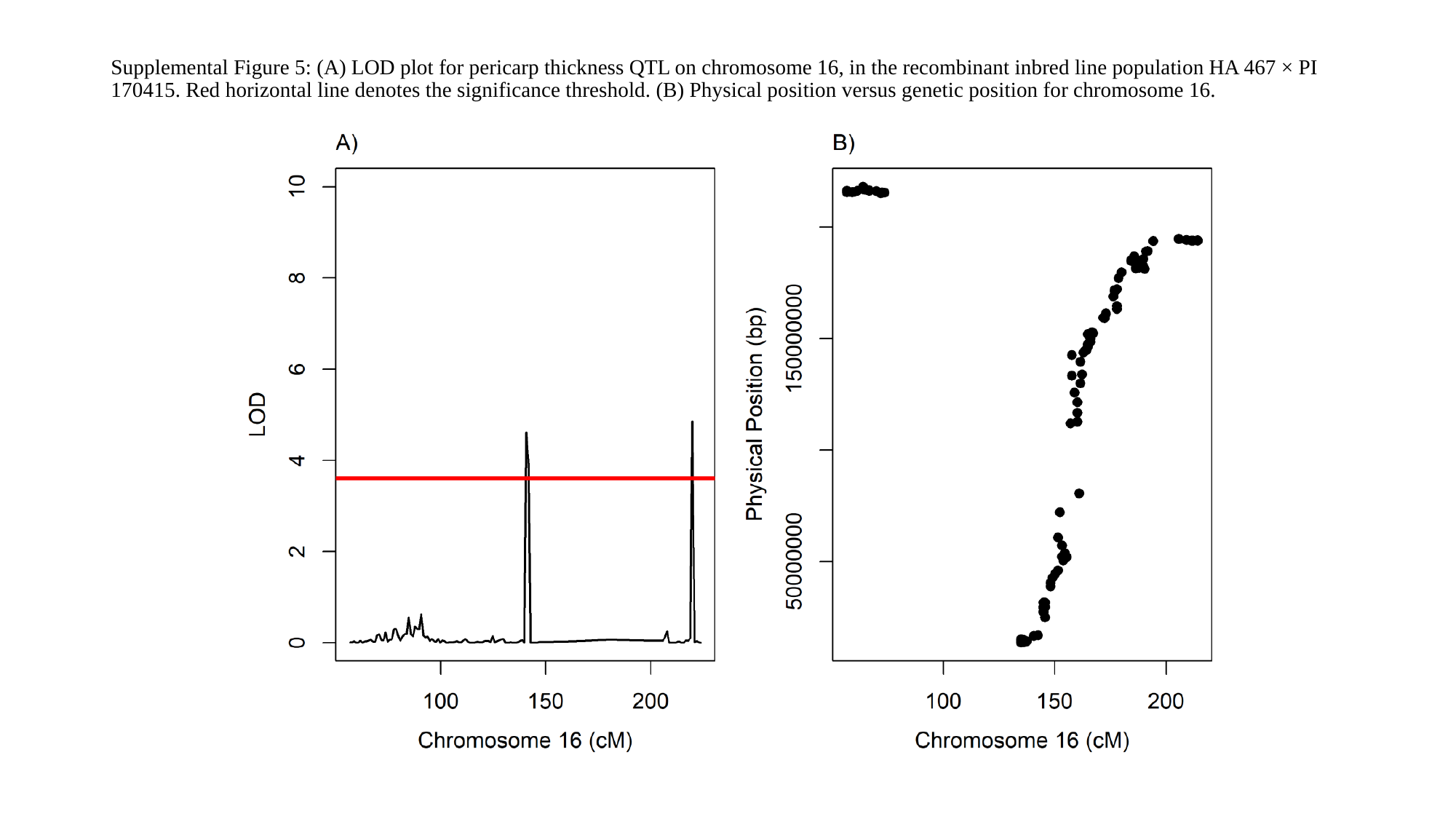

#### Supplemental Figure 5: (A) LOD plot for pericarp thickness QTL on chromosome 16, in the recombinant inbred line population HA 467 × PI 170415. Red horizontal line denotes the significance threshold. (B) Physical position versus genetic position for chromosome 16.
